# FastSKAT: Sequence kernel association tests for very large sets of markers

**DOI:** 10.1101/085639

**Authors:** Thomas Lumley, Jennifer Brody, Gina Peloso, Alanna Morrison, Kenneth Rice

## Abstract

The Sequence Kernel Association Test (SKAT) is widely used to test for associations between a phenotype and a set of genetic variants, that are usually rare. Evaluating tail probabilities or quantiles of the null distribution for SKAT requires computing the eigenvalues of a matrix related to the genotype covariance between markers. Extracting the full set of eigenvalues of this matrix (an *n* × *n* matrix, for *n* subjects) has computational complexity proportional to *n*^3^. As SKAT is often used when *n* > 10^4^, this step becomes a major bottleneck in its use in practice. We therefore propose fastSKAT, a new computationally-inexpensive but accurate approximations to the tail probabilities, in which the *k* largest eigenvalues of a weighted genotype covariance matrix or the largest singular values of a weighted genotype matrix are extracted, and a single term based on the Satterthwaite approximation is used for the remaining eigenvalues. While the method is not particularly sensitive to the choice of k, we also describe how to choose its value, and show how fastSKAT can automatically alert users to the rare cases where the choice may affect results. As well as providing faster implementation of SKAT, the new method also enables entirely new applications of SKAT, that were not possible before; we give examples grouping variants by topologically assisted domains, and comparing chromosome-wide association by class of histone marker.

## Introduction

The Sequence Kernel Association Test (SKAT) (Wu et al., 2011) is widely-used to test for associations between a phenotype and a set of genetic variants. SKAT provides a pooled test of multiple rare variants, and so is distinct from methods such as BOLT (Loh et al., 2015) and EM-MAX (Kang et al., 2010) that provide large numbers of single-variant associations. SKAT, a form of variance-components test, is particularly popular in analysis of rarer variants (e.g. minor allele frequency<0.05) where variant-by-variant analyses would have poor power, due to the large multiple-testing burden. Compared to tests that combine a region’s genotypes into a single “bur-den, SKAT retains power better when the true associations are heterogeneous (Lee et al., 2014). SKAT was initially developed for linear regression analyses in unrelated samples, but due to its popularity has been extended to analysis with logistic regression, proportional hazards regression and related subjects (Wu et al., 2010, 2015; Lee et al., 2012b,a; Chen et al., 2013a, 2014), among others.

With the advent of large-scale whole-genome sequencing (WGS) data (Cirulli and Goldstein, 2010), SKAT has been suggested for use testing for phenotype-genotype association, either in pre-specified regions of interest (Sung et al., 2014) or in ‘sliding windows’ (Morrison et al., 2013) across the entire genome. However, the computational burden of current SKAT code limits the number of variants that a single analysis can use. The rate-limiting step is calculating the eigenvalues of the covariance matrix of the genotypes, or those of a closely-related matrix—the computational complexity of which scales with the cube of the number of variants (*m*) or the number of subjects (*n*), whichever is smaller (Golub and Van Loan, 1996). In current large-scale WGS studies where *n* may currently be 20, 000 or more, this means a SKAT analysis of *m*=10,000 variants requires a million times more computing resources than one with *m*=100 variants. Using faster implementations of standard algorithms will not solve this problem; the matrix code used (e.g. LAPACK Anderson et al. (1999) or BLAS (Blackford et al., 2002)) is already heavily optimized (e.g. the high-performance version of level-3 BLAS (Goto and Van De Geijn, 2008)). Cluster-computing approaches—i.e. using a million times more processors—are prohibitively expensive, as well as being currently unavailable in pre-packaged software. But analyzing sets of *m*=10,000 variants is not unrealistic for WGS samples, where hundreds of millions of variants are observed. Moreover, sample sizes for WGS studies are increasing rapidly, so even when *n* is smaller than *m*, analyses will be hampered by this computational bottleneck. Novel statistical methods that avoid this problem are needed now.

To address this problem, we propose the *fastSKAT* method, that provides a highly-accurate approximation of SKAT‘s *p*-value with orders of magnitude smaller computational burden than current methods. FastSKAT achieves this in part using recent advances in random matrix theory (Halko et al., 2011; Tropp, 2011) that compute just the leading eigenvalues terms in the SKAT test. In this sense it is similar to the work of Galinsky et al. (2016), who similarly speed up principal components analysis. But fastSKAT also uses a form of Satterthwaite approximation to obtain *p*-values, that computes the most important terms exactly and uses a simple and expedient approximation for the remaining terms. Where standard SKAT’s computational burden grows with the cube of *m* (or *n* if smaller), the random-projection version grows only with the square of m. As well as the obvious speedup for SKAT analysis with large *m*, fastSKAT makes SKAT analyses possible for far larger sets of variants than are currently feasible. This means it can be used for entirely new forms of analysis, of which we give two examples. First, we perform SKAT analyses that group variants by topologically assisted domains (TADs), regions whose high conservation across species makes them a compelling way to cluster potential signals, but whose large size (10,000–20,000 rare variants) makes current SKAT analysis impractical. We also implement SKAT chromosome-wide, assessing the association of outcomes with different histone regulatory marks, and so learning which classes of variants to prioritize for subsequent localized inference.

## Material and Methods

### Overview of Methods

We first describe the SKAT (Wu et al., 2011) approach; its formulation, test statistic, and the null sampling distribution for that statistic. Our major focus is the computational burden of evaluating all eigenvalues of the genotype matrix, which is a limiting factor in SKAT analysis of WGS data. We then describe the fastSKAT method, which computes just the leading eigenvalues terms in the SKAT test using random projection methods (Halko et al., 2011; Tropp, 2011) and related tools. Instead of evaluating every eigenvalue, these methods focus on evaluating just the leading (i.e. largest) eigenvalues, by examining the eigensystem of a random projection of the original matrix. The projection is low-dimension, and so its eigenvalues can be computed quickly, but as the leading eigenvalues are those best-preserved under projection, they are the ones a random projection is most likely to be informative about. Averaging the process over different randomly-chosen projections, the error in the approximation of these leading eigenvalues quickly becomes negligible. We describe various options within fastSKAT that can further optimize its speed and accuracy in different applied settings. Finally, we also describe the settings for simulation-based evaluation of fastSKAT, and its practical application.

An R package implementing fastSKAT and providing further examples of its use is available from https://github.com/tslumley/bigQF.

### SKAT

SKAT (Wu et al., 2011) can be derived as a score test of the null distribution that all variants have no association versus the alternative that their effects follow a Gaussian distribution. In the absence of covariates, it uses an *m* × *n* genotype matrix *G*, where *m* is the number of variants and *n* is the number of individuals (e.g., each row is a variant, each column is a sample). Each entry in this matrix takes its values from {0,1,2} indicating the count of variant alleles for a sample at a variant. From this matrix we generate *g*, the *m*-vector of observed minor allele frequencies *g_j_* for variant *j* = 1,2,…, *m*. SKAT also uses phenotypes vector *Y* (of length *n*) and corresponding vector *µ* of predicted means, either a constant vector or taken from a linear model that uses adjustment variables denoted *X*. The SKAT test statistic is

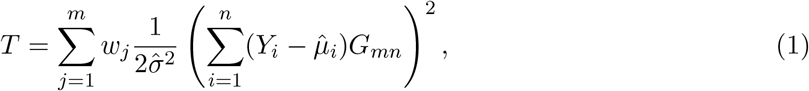

where 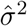 is the sample variance of 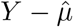, and the weights denoted by *m*-vector *w* are a function of *g* and possibly of annotation data; in the default version of SKAT, the ‘Wu weights’ use *w_j_* = 25(1 - *g_j_*)24.

Under a classical linear model (or just under mild regularity conditions, in large samples) and under the null hypothesis of no genetic effects, the distribution of *T* can be described as a sum of independent χ^2^ variables, specifically

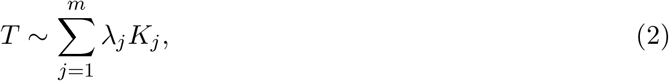

where the λ_*j*_ are the eigenvalues of an *m* x *m* matrix (described below) that is closely related to the covariance of the genotypes, and are ordered from largest to smallest. Given these λ_*j*_, to efficiently compute tail probabilities and quantiles of *T*, we can use the exact methods of Davies (1980); Farebrother (1984), or approximate them with high accuracy using saddlepoint methods (Kuonen, 1999), with computation proportional to *m* in all cases.

An important preliminary step is the calculation of the λ_*j*_. One succinct way to write them is as the eigenvalues of 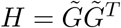, where

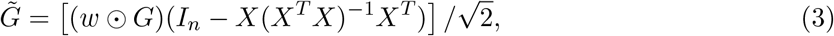

in which *I_n_* denotes an identity matrix, 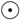 denotes the Hadamard (elementwise) product, and *X* is the usual ‘design’ matrix of adjustment covariates, including an intercept. Equation (3) is not a computational formula - for example, the projection orthogonal to the range of *X* given by the matrix (*I* — *X*(*X^T^X*)^−1^*X^T^*) can be computed more efficiently using the *QR* decomposition of *X*. (The computation of the elements of *H* and the relevant decomposition of 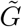 is further discussed in Appendix 1.) But regardless of how matrix *H* is evaluated, calculating all the eigenvalues of an *m* x *m* matrix in general has computational complexity proportional to *m*^3^. If *n* ≪ *m* we can instead work with the *n* x *n* matrix 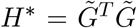 (Price et al., 2006) and compute the *n* non-zero eigenvalues in O(*n*^3^) time, but for WGS applications both *n* and *m* tend to be large.

### fastSKAT: Satterthwaite approximations

We propose to compute *p*-values for SKAT tests using a form of Satterthwaite approximation (Lumley, 2011). The basic Satterthwaite approach approximates the reference distribution of *T* by a single scaled χ^2^ distribution, i.e.

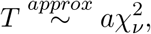

with scaling factor *a* and degrees of freedom *ν* selected by moment-matching arguments, so that

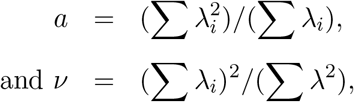

where the summation is over all *m* or *n* eigenvalues. Unfortunately, for SKAT tests the basic Satterthwaite approximation tends to be anticonservative (see e.g. Schifano et al. (2012)), often misstating *p*-values in the vicinity of 10^−6^ by an order of magnitude. Details are given in Appendix 2, but briefly the 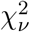 distribution, for any *ν*, has tails that are too light compared to the weighted sum of 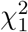 distributions given in (2). The basic Satterthwaite approximation is sufficiently accurate for filtering—if it gives a *p*-value larger than, say, 10^−3^ in a genome-wide scan no further computation is needed—but this method alone cannot be sufficiently accurate for final decision-making.

To improve on the basic Satterthwaite approach, we propose instead using

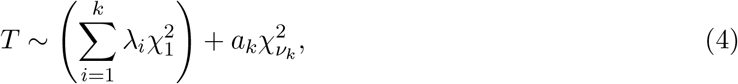

where λ_i_,…, λ_k_ are the largest *k* eigenvalues of *H*. Moment-matching arguments for the scaling and degrees of freedom in the ‘remainder’ term give

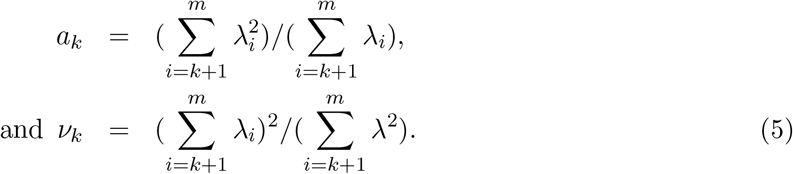

Since 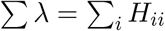 and 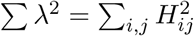 this takes time proportional to *m*^2^, with a small constant of proportionality.

In the Results section we compare fastSKAT’s approximation to a simpler low-rank approximation with no remainder term. We also consider the four-moment approximation of Liu et al. (2009), which is one option in the `SKAT` package (Lee et al., 2016). This estimator does not require individual eigenvalues, but does require the trace of powers of *H* up to *H*^4^, and calculating these matrices explicitly is as expensive as extracting the full set of eigenvalues.

### fastSKAT: fast eigenvalue calculations

For the problem of calculating λ_1_,…, λ_k_ in (4) we propose using a random projection using the Subsampled Random Hadamard Transform (Tropp, 2011), which multiplies each row of *H* by a random sign, applies the Fast Hadamard Transform to the matrix, and samples *k*+*p* rows at random from the result. Let Ω be the matrix corresponding to this linear transformation. When working with *H* we use the *QR* decomposition of (Ω*H*)*^T^* to produce an orthonormal matrix *Q* and compute the eigenvalue decomposition of *QHQ^T^*. When working with 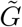 we similarly compute *Q* from Ω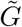 and then take the singular value decomposition of Q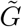. The *k* largest eigenvalues of *QHQ^T^* and singular values of Q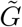 will be good approximations to those of *H* and 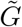 respectively.

Still better approximations are available by working (implicitly) with the matrix (*HH^T^*)^q^*H*. After forming *Q* = *Q*_0_ from (Ω*H*)*^T^*, we compute *H^T^Q* and do a *QR* decomposition to extract 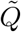, then form H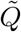 and do a *QR* decomposition to extract an improved *Q*_1_, or by iterating the procedure *q* times, *Q_q_*(Halko et al., 2011, Algorithm 4.4). Our implementation defaults to *q* = 3. Each iteration takes O(*nmk*) operations, so the total time complexity is proportional to *q* + 1.

Because of the use of the Fast Hadamard Transform the construction of *H*Ω takes only O(*m*^2^ log(*m*)) operations, and the entire approximate singular value decomposition (SVD) only *O*(*m*^2^ log(*m*) + *mk*^2^). Working with 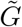, the complexity is *O*(*nm* log *n* + (*m* + *n*)*k*^2^), and the iterated improvement changes the leading term to O(*nmk*), much smaller than the min(*m*^3^, *n*^3^) needed for a complete eigendecomposition.

For situations where *H* is too large to fit in memory, Halko et al. (2011, Section 5.5) describe single-pass versions of the algorithm; we do not consider these here but they may become useful in the future.

Another approach for estimating eigenvalues is Lanczos-type algorithms, which generalize the power algorithm (Golub and Van Loan, 1996). Briefly, this uses an arbitrary starting vector *υ*, transforming the vectors *Aυ*, *A*^2^*υ*, *A*^3^,…, *A^k^υ* into an orthogonal basis in which the matrix *A* will be tridiagonal and its eigenvalues easily computed. The basic power algorithm is prohibitively inaccurate with finite-precision arithmetic, but a variety of modified algorithms have been constructed that give the *k* largest eigenvalues accurately, though with more than *k* matrix multiplications. Here, for comparison with fastSKAT’s Satterthwaite approach, we use the nu-TRLAN implementation of the thick restarted Lanczos algorithm (Yamazaki et al., 2010), as provided in the svd package for R (Korobeynikov et al., 2016).

### fastSKAT: fast trace estimators

FastSKAT uses Satterthwaite approximation to calculate the ‘remainder’ term in the distribution of *T*. Write *F* = *H*—*QQ^T^H* for the remainder when the low-rank approximation is subtracted from *H*. The approximation requires the trace of *F* and of *F^T^ F* and their low-rank approximations.

The former is computationally straightforward, as the diagonal of *H* can be computed in *O*(*mn*) time, followed by subtracting the *k* leading eigenvalues (in *O* (*k*) time). The most efficient way to compute the latter exactly is to construct *H* and use 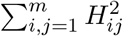, but this would take *mn*^2^ time. Instead, we approximate it by a version of Hutchinson’s randomized trace estimator (Hutchinson, 1990).

Specifically, let *υ_i_* be *m* random *m*-vectors with 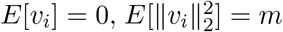 and 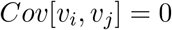. Define 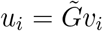 and 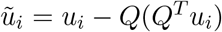, so that 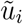 is the projection of *υ_i_* orthogonal to *Q*. An estimator of the trace of the remainder of HT *H* using only multiplications by 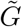 and 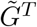 is

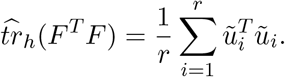

The estimator takes *O*(*rmn*) time to compute, and by the central limit theorem has relative error *O*_*p*_(*r*^−1/2^).

### fastSKAT: stabilizing ratio estimates

FastSKAT’s Satterthwaite approximation, given in (5), uses the ratio of terms that are traces. We can therefore increase their accuracy by calculating them using a survey ratio estimator (Fuller, 2011, Section 2.1). Using a randomized trace estimator with the same random *υ_i_* to obtain an estimate 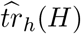, since 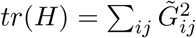 is available exactly, we can compute

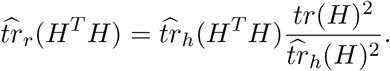

As the Monte Carlo errors in 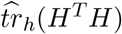 and 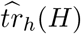 are correlated, the ratio estimator has increased accuracy.

### fastSKAT: ‘matrix-free’ methods for unrelated individuals

The matrices *H* and 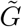 from (3) will not typically be sparse, but in unrelated individuals 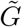 is the product of a sparse matrix and a projection orthogonal to a matrix of low rank. The projection is on to residuals for the adjustment model, and so for *p* adjustment variables can be computed in *O*(*np*^2^) time from the *QR* decomposition of *X* that was computed to fit the adjustment model.

In order to take advantage of this representation we can replace the implicit random matrix Ω (from the material above on fast eigenvalue calculations) by an explicit *n* x (*k* + *p*) matrix of random standard Normal variables. Forming *H*Ω now takes *k* + *p* matrix-vector multiplications by H, followed by the same *QR* decomposition and eigenvalue decomposition as before. We also need to replace the implicit random matrix in the trace estimator by an explicit matrix, and again we use random standard Normal variables. Finally, when there is an adjustment model, use the randomized trace estimator for both 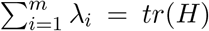 and 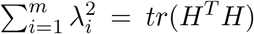. The cost of computing *H*Ω is now proportional to *mnkα*, where *α* is the fraction of non-zero entries in *G*. We call this approach ‘matrix-free’; while not described in detail here a similar approach can be used with the Lanczos-type algorithms. Both are supported by our R package.

### fastSKAT: special methods for family data

Adaptations of SKAT’s test statistic (1) and its distribution (2) are available for use with family data, under a polygenic model for residual phenotype variance (Chen et al., 2013b). Let Φ be the *n* x *n* kinship matrix and 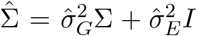 be the phenotype covariance matrix, estimated under the null of no SNP effects. Replacing 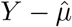 in (1) by 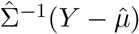, *G* is then defined by

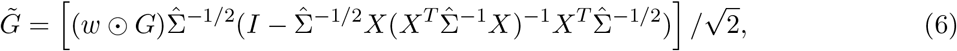

and the distribution of *T* in (1) holds. Analogous results for a binary phenotype and logistic adjustment model are provided by Wu et al. (2010).

When 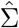 is based on expected kinship (pedigree) rather than on observed identity-by-state it is often sparse, with a sparse Cholesky factorization. The sparseness allows 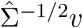, for a vector *υ*, to be computed in *O*(*nf*^2^) time where f is the size of the largest pedigree - with sharper bounds possible using the distribution of pedigree sizes. In the setting of (6) this means that, if the size of the largest pedigree is bounded, the time complexity for large *m*, *n* only exceeds that for unrelated individuals by only a constant factor, when *m* and *n* are of the same order.

### Data analysis settings and simulation framework

#### Comparison with standard SKAT

We performed standard SKAT and fastSKAT analysis of data from CHARGE-S, an early, small-scale WGS study in unrelateds (Lin et al., 2014) conducted by investigators from the CHARGE consortium (Psaty et al., 2009). The disease outcome is low density lipoprotein (LDL), adjusted to account for lipid-lowering medication use, and analyses residualized out effects of age, cohort, study site and 5 principal components of ancestry (Morrison et al., 2013) before inverse-Normal transformation. Tests were performed on regions of typical size centered on known genes (transcript ±50Kb). The fastSKAT method used *k*=100 eigenvalues and *B*=600 random projections.

#### Comparing versions of fastSKAT

To assess the performance of the fastSKAT approximation (4) to other SKAT implementations, and to assess the speed and accuracy of the various versions of fastSKAT, we used simulated human genome sequence data. The data was generated using the Markov Coalescent Simulator (Chen et al., 2009), fixing *n* and choosing the sequence length to given *m* ≈ *n*. We discarded variants with minor allele frequency over 5%. The resulting genotype matrix has about 98% zero entries.

We perform fastSKAT using the proposed approximation and a low-rank approximation with no remainder term. To provide a fair assessment of each approximation’s performance, this comparison is based on a full eigendecomposition of *H* and so does not include any Monte Carlo error from the randomized trace estimator or similar algorithms.

Unless specified otherwise, we evaluated the approximations at the point where the Satterthwaite inequality gave a *p*-value of 10^−6^, for a continuous phenotype, unrelated samples, and no adjustment variables.

Simulations were conducted on an iMac with 8GB memory and a 3.4GHz Intel Core i5 processor, using R 3.2.1(R Core Team, 2016).

#### Data analysis

To illustrate fastSKAT analysis of much larger regions than standard SKAT, we provide results from analysis of the CHARGE-S LDL data as described above, but aggregating rare variants by “topologically associated domains (TADs). TADs, typically 1Mb wide, are regions that mark higher order chromatin interaction (Yao et al., 2015) and are found across the human genome. In this example we use Human ES Cell TADs (Dixon et al., 2012), which in this setting typically contain 10,000-20,000 rare variants.

We also used fastSKAT in chromosome-wide analyses, to examine the relative contribution of rare (below 1% MAF) variants that fall within regulatory marks of six histones annotated in adult liver and within 500Kb of known lipid loci. This was done for the same CHARGE-S LDL data as above, but now producing a single SKAT test for each chromosome, for each of the six histones. Random-selected sets of the same number of SNPs drawn from the same regions, chromosome-wide, were tested for comparison.

## Results

### Comparison with standard SKAT

Figure 1 shows a comparison of fastSKAT and standard SKAT, in the setting of the CHARGE-S study – where standard SKAT is computationally feasible. The analysis gave 28912 tests overall, for which standard SKAT took approximately 190 CPU hours, while fastSKAT took only 10 CPU hours. As shown, the agreement between methods is perfect for all practical purposes. As a positive control, we note that the eight gene regions where SKAT gave significant results (after Bonferroni correction) all surround the APOE locus. The agreement between the methods means that their statistical properties (e.g. Type I error rate, power) can be considered equivalent; for statistical properties of fastSKAT we therefore refer to the literature on standard SKAT (Lee et al., 2014).

**Figure 1:**
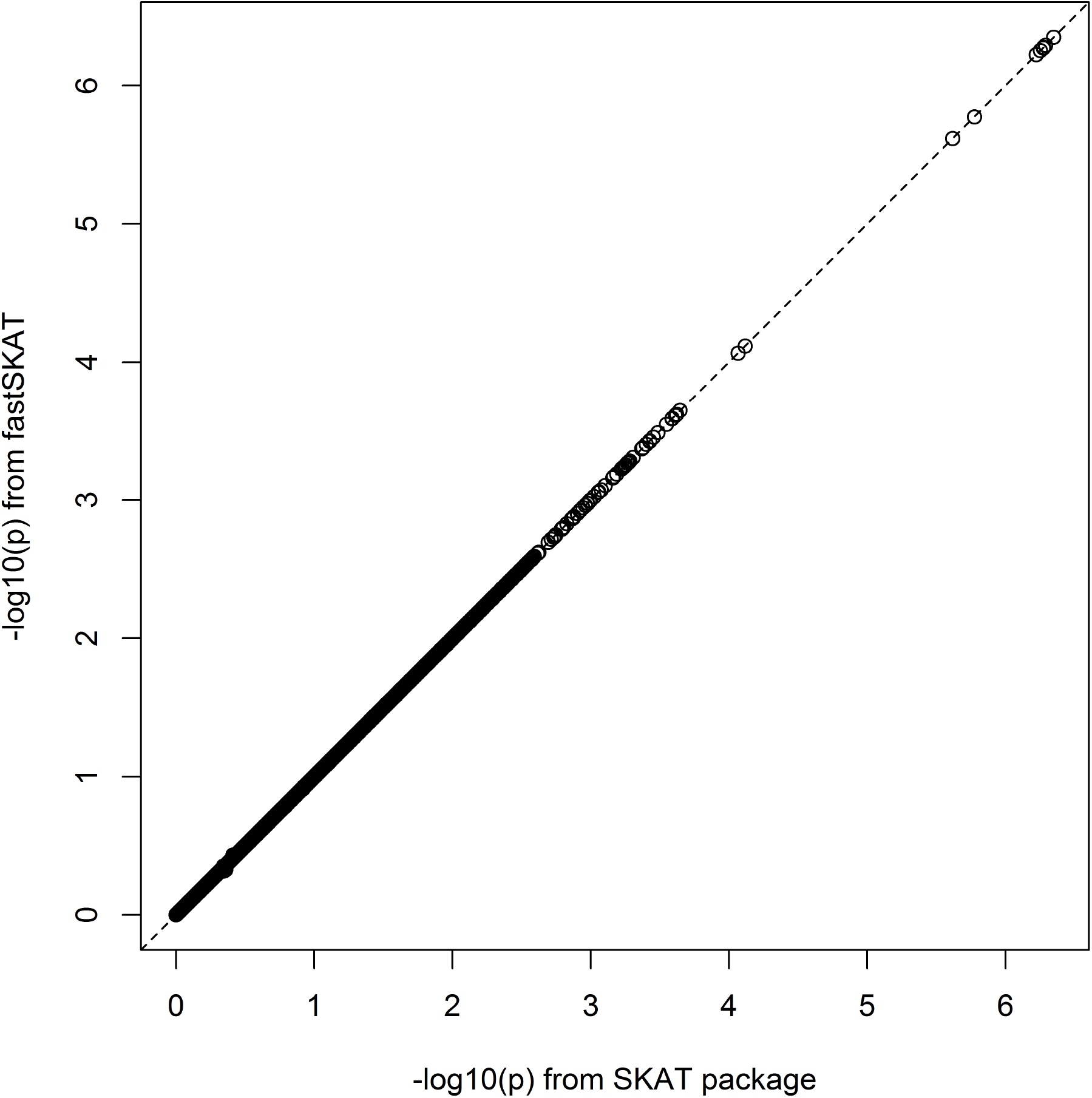
Scatterplot of fastSKAT versus standard SKAT *p*-values (-log_10_ scale) for pilot LDL analysis using CHARGE-S data.

### Comparing versions of fastSKAT

As Figure 2 shows, the approximations using a remainder term perform much better with much lower values of *k* than an approximation with no remainder term. Figures 3 and 4 show the total computation time and relative error in the log_10_ *p*-value (log_10_ *p*_fastSKAT_ - log_10_ *p*_SKAT_) for various approximations when *H* is assumed to be already available (and hence not included in the timing calculations), and when computing *H* is included. It shows that ‘matrix-free’ fastSKAT is much faster than SKAT, with little approximation error, and that when *H* is not already available, computing it is a computational bottleneck. The moment-based approximations have substantially greater approximation error at these extreme *p*-values than fastSKAT, and are not importantly faster.

Figures 5 and 6 are similar, but with *M* = 10^4^ and *n* ≈ 7500. Initial experimentation showed that the 100-eigenvalue approximation was not as good as with *m* = 5000, having typical error of 0.1 on the log_10_ *p* scale. These simulations used 200 eigenvalues, giving better approximation than 100 eigenvalues for *m* = 5000 had done. To put the magnitude of these errors in perspective, the random fluctuation in signals is much larger. For example, with a signal where a comparison to 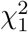 gives median *p*-value= 10^−6^, we would expect — log_10_ *p* to vary between 2.47 and 11.14 in 95% of repeat experiments.

Figure 7 show the impact of parameter choice on approximation accuracy, where the full eigende-composition is not available. The left panel is for the algorithms using H, the right panel is for the algorithms using 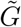. Since 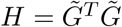, an increase of 1 in *q* in the left panel has the same effect as an increase of 2 in the right panel. Clearly *q* ≥ 1 is desirable for stochastic SVD of *H* and *q* ≥ 2 for 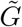. We chose *q* = 1 and 100 eigenvalues when using *H*; *q* = 3 and 100 eigenvalues when using 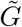.

**Figure 2:**
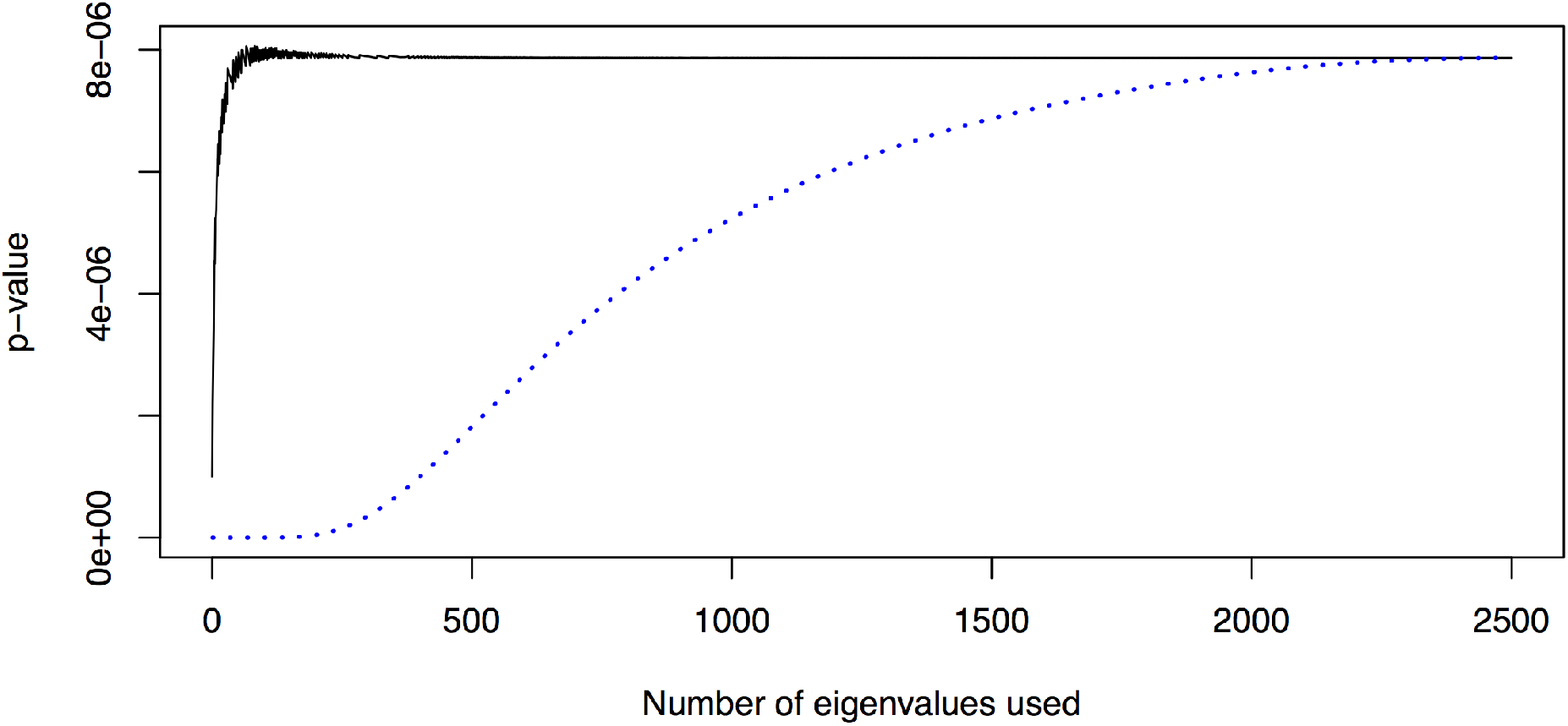
Comparing a rank-k approximation to *H* (dotted line) to the proposed approximation (solid line) for a single simulated dataset with *m* = 4028, *n* = 5000.

**Figure 3:**
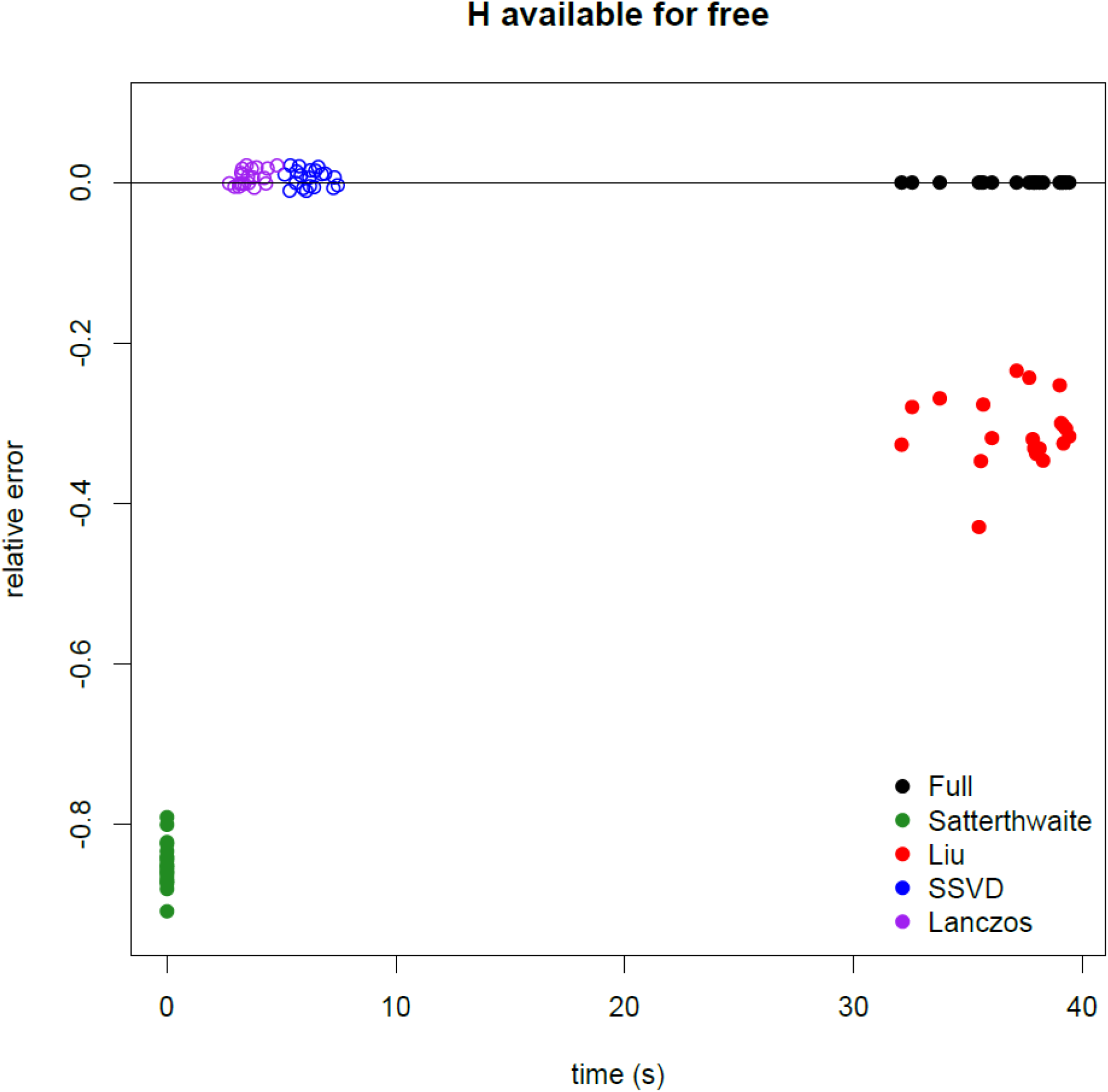
Computation time and error for a *p*-value near 10^−6^ when the square matrix 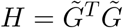 is already available (*n* = 5000, *m* ≈ 4000). Solid points are deterministic approximations, hollow points are based on sampling. Lanczos and SSVD use 100 eigenvalues.

**Figure 4:**
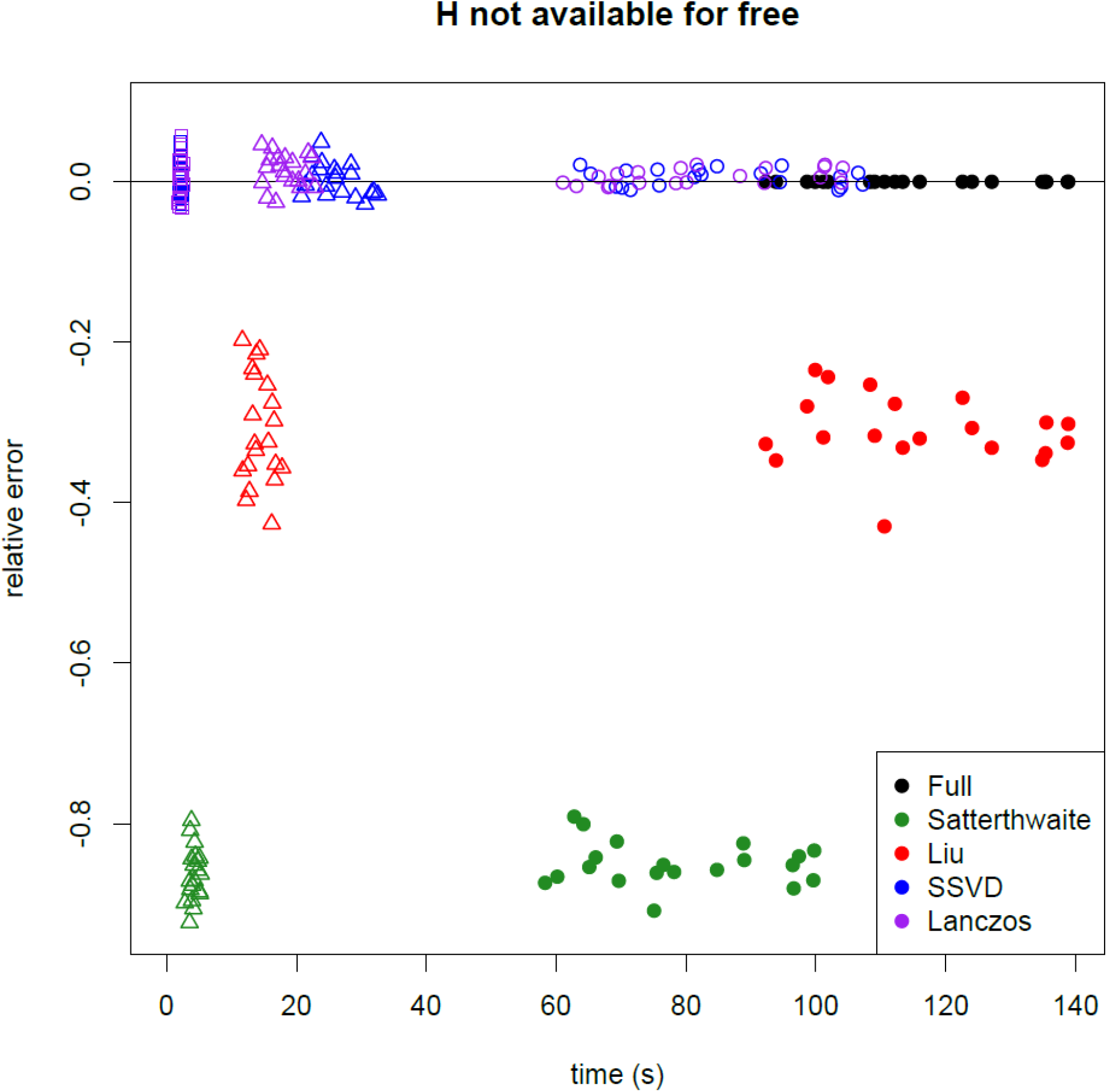
Computation time and error for a *p*-value near 10^−6^ including the cost of computing *H* when needed (*n* = 5000, *m* ≈ 4000). Lanczos and SSVD use 100 eigenvalues. Solid points are deterministic approximations, circles are based on sampling, triangles based on 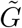, and squares based on sparse *G*.

**Figure 5:**
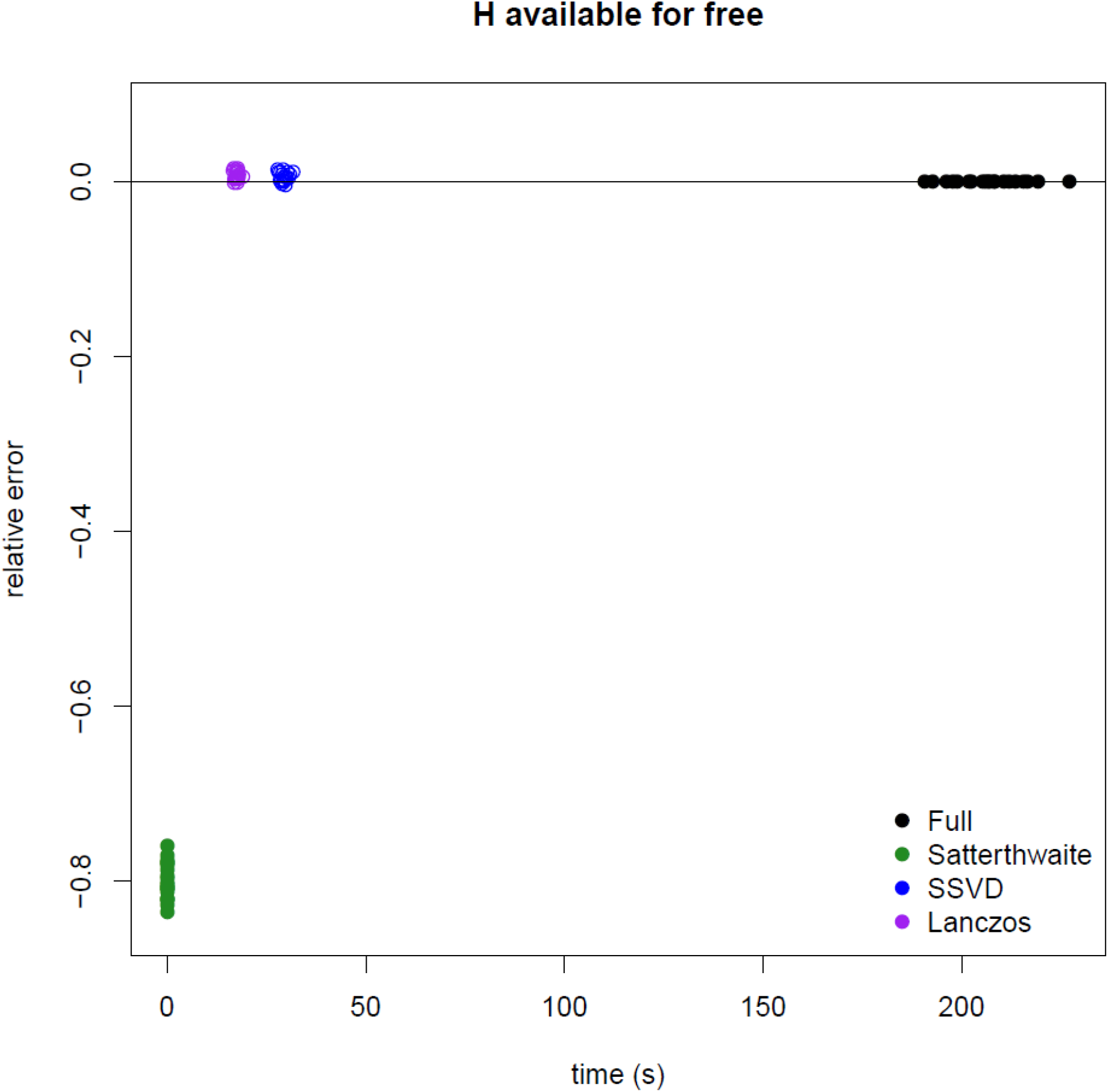
Computation time and error for a *p*-value near 10^−6^ when the square matrix 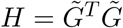 is already available (*n* = 10^4^, *m* ≈ 7500). Solid points are deterministic approximations, hollow points are based on sampling. Lanczos and SSVD use 200 eigenvalues.

**Figure 6:**
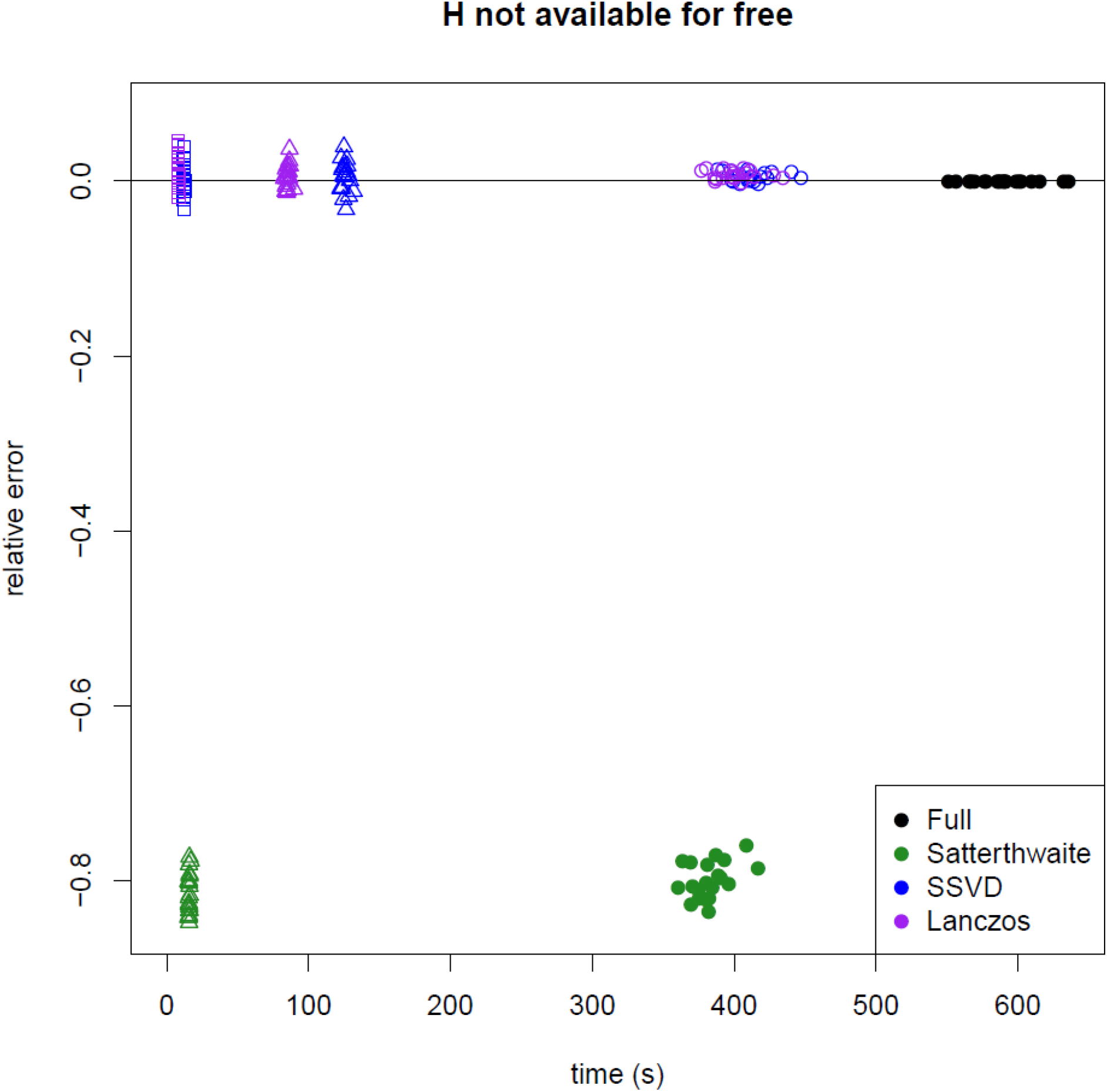
Computation time and error for a *p*-value near 10^−6^ including the cost of computing *H* when needed (*n* = 10^4^, *m* ≈ 7500). Lanczos and SSVD use 200 eigenvalues. Solid points are deterministic approximations, circles are based on sampling, triangles based on 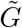, and squares based on sparse *G*.

**Figure 7:**
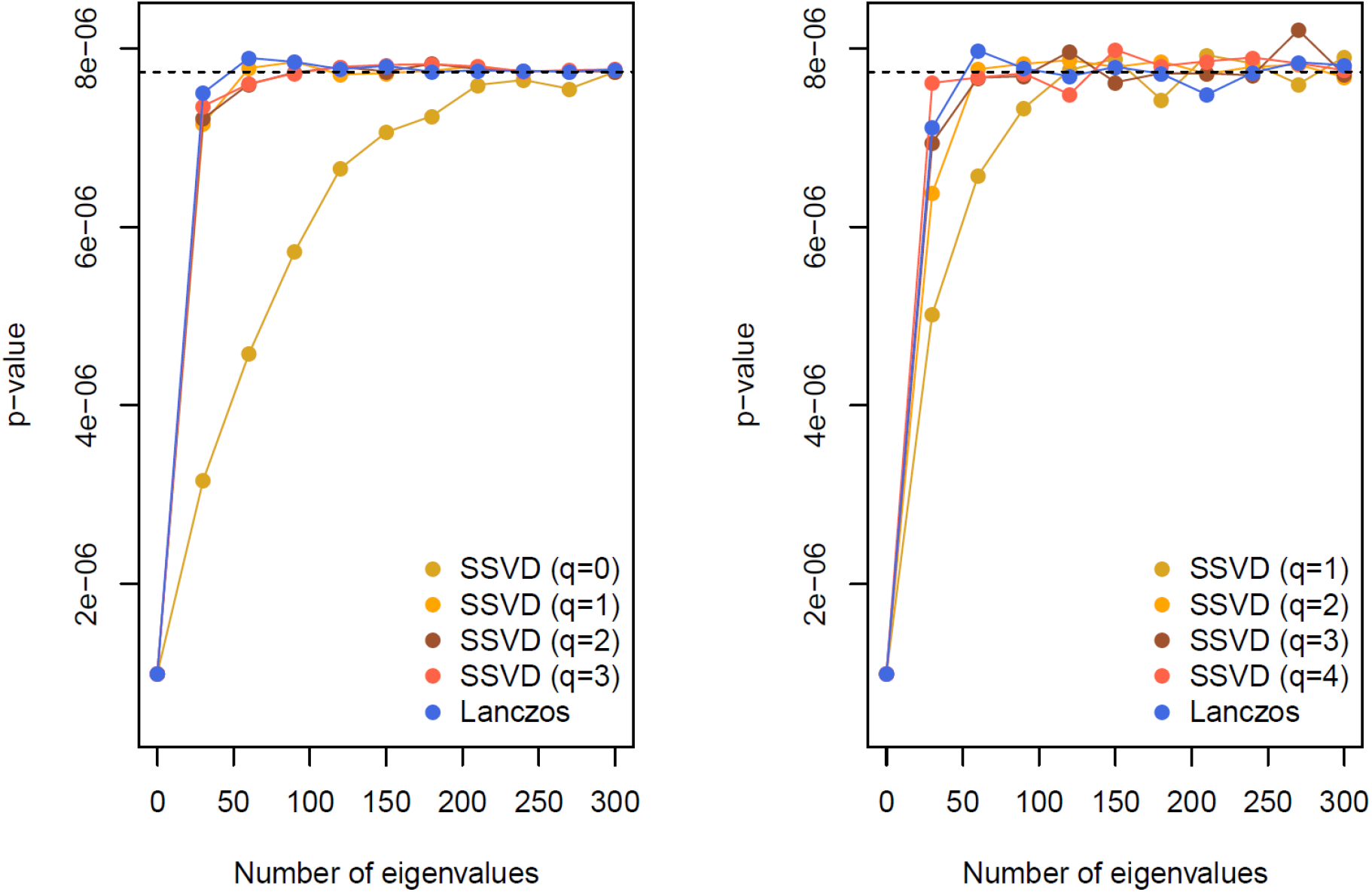
Dependence of accuracy on number of eigenvalues used and on the iteration parameter *q*, with *n* = 5000, *m* = 4151. Left panel: using *H*, right panel: using 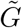. A difference of 1 in *q* in the left panel is equivalent to a difference of 2 in the right panel. The dashed line indicates the *p*-value based on a full eigendecomposition.

### Comparing choice of k

As Figures 2 and 7 show, the accuracy of the approximation is not sensitive to *k* as long as *k* is large enough. As a simple criterion, we recommend checking if *k* and 2*k* give similar results in a selection of genes; if so, *k* can be assumed large enough to give the correct results. Our implementation defaults to *k* = 100 eigenvectors and *r* = 500 random projections.

The choice of *k* is more important when *M* or *N* is very small. Although there is no direct benefit to using the approximation for min(*m*, *n*) < 200, there is an advantage to having a consistent computational pipeline, so we investigated the small example (*n* = 2000, *m* = 67) provided with the SKAT package(Lee et al., 2016). In this example using the full eigendecomposition gives a *p*-value of 0.01875. The leading-eigenvalue approximation based on *H* (and otherwise using the default settings) gives *p*-values with an error of less than 1 in the third decimal place, for all *k* from 5 to 60. Using 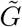 and the randomized trace estimator, the accuracy is at a similar level; it appears safe to use fastSKAT even when it is not necessary for computation.

**Figure 8:**
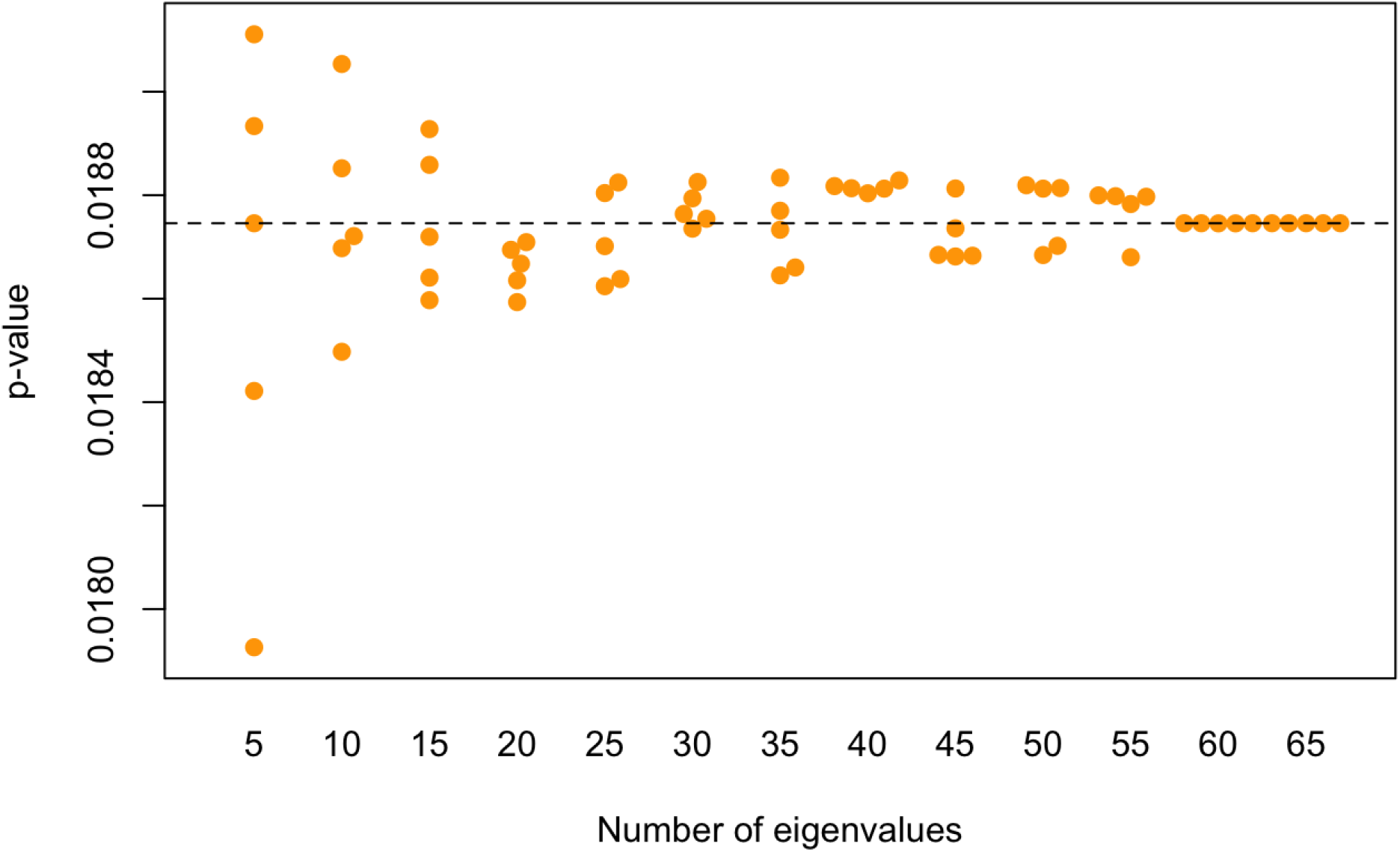
Small-sample behavior of matrix-free fastSKAT: *k* = 1, 2, 3, …, 65

### Data analysis: aggregating by TADs

The results are shown in Figure 9, and are in close accordance with what would expected under the null, as might be expected from this small-scale WGS study. After correction for the number of regions (n=2977) no region’s association, as measured by SKAT, was statistically significant. However, the top TAD signal (p=8.2e-5) contains the known lipid gene APOE, a well-known LDL gene.

In terms of computation, the entire fastSKAT run took 16 CPU hours, reduced to 15 minutes by parallelization. Using standard SKAT it would have taken approximately 260 CPU days, with corresponding greater costs even if parallelized.

**Figure 9:**
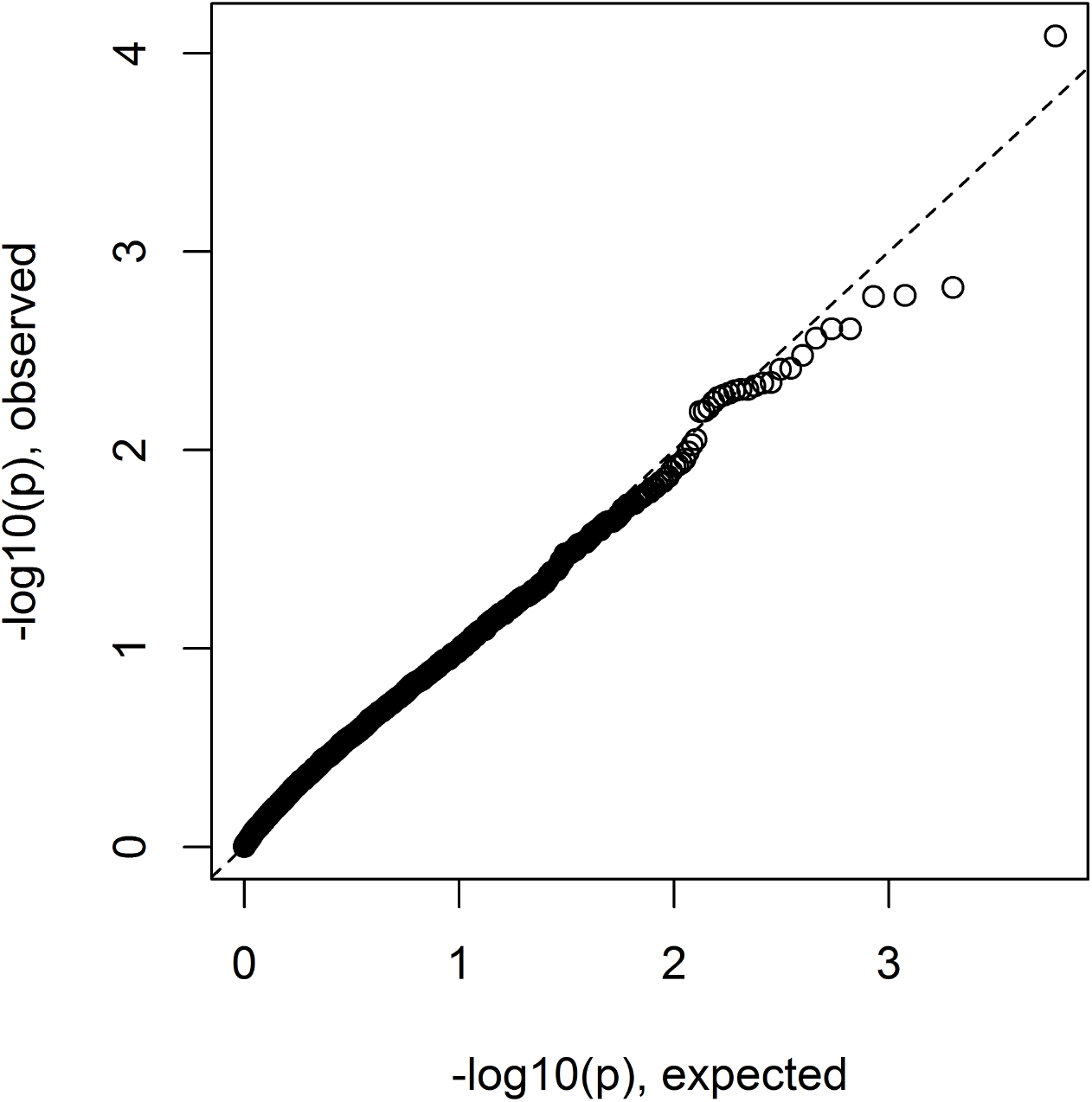
Genome-wide results for LDL association with rare variants aggregated by TADs.

### Data analysis: chromosome-wide association by histone class

The results are shown in Figure 10. Computation of the whole analysis took approximately 1 CPU hour. Two chromosome-wide aggregations of histone variants were significant after correcting for the number of chromosomes and histones, variants within H3K4me1 on chromosome 19 (p=1.3e-4) and H3K36me3 (p=2.0e5). Random variants from the same regions were associated at 0.02 and p=0.08 respectively. No systematic differences in association strength were detected across histone marks. As well as suggesting that chromosome 19 is the most promising location for more detailed examination of the data, the results highlight the considerable strength of evidence brought by a priori knowledge – in this case histone-related variants over random selection of variants.

**Figure 10:**
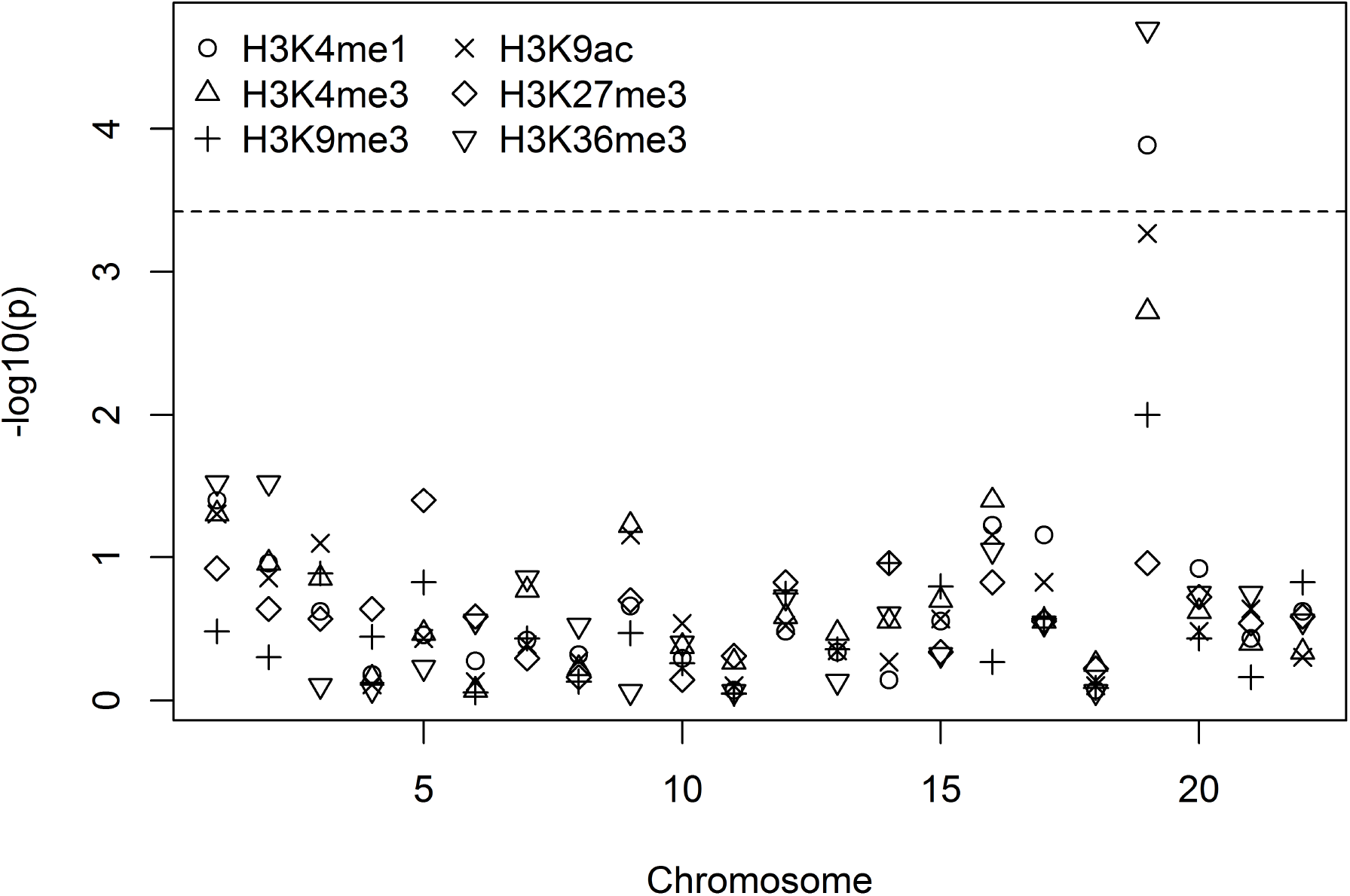
Chromosome-wide results for the autosomes, for SKAT association tests for all rare variants within regulatory markers of the six listed histones, and withing 500Kb of known loci. The dashed line indicates Bonferroni correction of level 0.05 for 22 autosomes × 6 histones.

## Discussion

We have provided the fastSKAT method, that implement the well-known SKAT test with large numbers of variants and/or large sample sizes. The fastSKAT method is approximate, but is sufficiently accurate for routine use with little to no tuning by the user. As well as providing notable speed increases for analyses that can currently be done, fastSKAT enables entirely new forms of analysis to be done, such as our aggregation by TADs or even across entire chromosomes.

A number of further refinements are available in special circumstances. If matrix *H* is already available (or the genotype covariance matrix, from which *H* can be computed rapidly), using it provides a substantial speedup, but computing *H* just to do the test is not efficient. Similar conclusions apply to computing *H* to examine population substructure or to look for duplicates, where similar leading-eigenvalue algorithms can be used. Taking advantage of the sparseness of *G* in genome sequence data can reduce computation time and memory use by a further large factor.

If there is uncertainty about the number of eigenvalues needed, e.g. for very much larger data sets than those considered here, we recommend increasing the number until the estimated *p*-value stabilizes. If *k* is too small, the results will change rapidly with increasing *k* and then stabilize for *k* ≪ (*m*,*n*). Such approaches are common in other approximation algorithms, e.g. quadrature (Fitzmaurice et al., 2012, Pg 411).

As seen here, the stochastic SVD and Lanczos algorithms have broadly similar performance. The stochastic algorithm is much easier to implement, but production-quality free implementations of Lanczos-type algorithms are available, making this less important. While not explored here, the stochastic SVD is easier to parallelize, which may be important in still-larger applications.

Finally, the Lanczos-type algorithms and randomized algorithms in linear algebra are not well known to applied statisticians. Based on our example, and the work of Galinsky et al. (2016) on principal components analysis, these methods have considerable potential and are likely to have many useful applications in analysis of large-scale genetic data.

## Acknowledgements

Funding support was provided by NHLBI grant U01HL137162 “From gene regions to whole chromosomes: scaling up association-finding for disease and omics outcomes in TOPMed. Fund-ing support for “Building on GWAS for NHLBI-diseases: the U.S. CHARGE Consortium was provided by the NIH through the American Recovery and Reinvestment Act of 2009 (ARRA) (5RC2HL102419). Data for “Building on GWAS for NHLBI-diseases: the U.S. CHARGE Con-sortium was provided by Eric Boerwinkle on behalf of the Atherosclerosis Risk in Communities (ARIC) Study, L. Adrienne Cupples, principal investigator for the Framingham Heart Study, and Bruce Psaty, principal investigator for the Cardiovascular Health Study. Sequencing was carried out at the Baylor Genome Center (U54 HG003273). The ARIC Study is carried out as a collaborative study supported by National Heart, Lung, and Blood Institute (NHLBI) contracts (HHSN 268201100005C, HHSN 268201100006C, HHSN 268201100007C, HHSN 268201100008C, HHSN 268201100009C, HHSN 2682011000010C, HHSN 2682011000011C, and HHSN 2682011000012C). The Framingham Heart Study is conducted and supported by the NHLBI in collaboration with Boston University (Contract No. N01-HC-25195), and its contract with Affymetrix, Inc., for genome-wide genotyping services (Contract No. N02-HL-6-4278), for quality control by Framingham Heart Study investigators using genotypes in the SNP Health Association Resource (SHARe) project. A portion of this research was conducted using the Linux Cluster for Genetic Analysis (LinGA-II) funded by the Robert Dawson Evans Endowment of the Department of Medicine at Boston University School of Medicine and Boston Medical Center. This CHS research was supported by NHLBI contracts HHSN 268200800007C, N01 HC85079, N01 HC85080, N01 HC85081, N01 HC85082, N01 HC85083, N01 HC85084, N01 HC85085, N01 HC85086, N01 HC35129, N01 HC15103, N01 HC55222, N01 HC75150, N01HC45133, HHSN 268201200036C and NHLBI grants HL080295, HL087652, HL105756 with additional contribution from NINDS. Additional support was provided through AG023629, AG15928, AG20098, and AG027058 from the NIA.

## Appendix 1 Complexity of calculating *H*

It remains to consider the complexity of calculating H. To simplify notation, we will assume *m* is not of smaller order than *n* and that *k* > logn; the latter assumption is likely to be true and the the former is no loss of generality as we can work with the transpose of *H\** instead of *H*. Computing *H* directly, if it is not needed for other reasons, has complexity proportional to *m*^2^*n*. While the proportionality constant is small, the task will eventually dominate the computational effort if *m* and *n* are both large. In that situation, it is possible to work with the singular value decomposition of 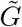 directly. Computing λ_1_,…, λ_k_ then takes *O*(*mnk* + *mk*^2^) time.

## Appendix 2 Theoretical behavior in the right tail

The anti-conservatism of the Satterthwaite approximation in the extreme tail is a general phenomenon, as can be proved using Theorem 3.1 of Berman et al. (1992) on tails of convolutions. Suppose two independent random variables have density functions *f*(*x*) and *g*(*x*) with exponential tails, in the sense that the limits

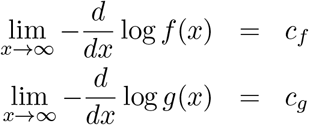

exist and are finite and non-zero. The theorem states that if *c_f_* < *c_g_* the sum of the variables also has a density *h*(*x*) with an exponential tail, and lim_x→∞_ *f*(*x*)*/h*(*x*) exists and is finite and non-zero, and consequently lim _x→∞_ *g*(*x*)*/h*(*x*) *=* 0.

Multiples of chi-squared densities have exponential tails, and if *f* is the density of 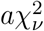, the tail rate *c_f_* = (2*a*)^−1^ depends only on the multiplier, not on the degrees of freedom. Thus, the extreme tail of the density of *T* is exponential with rate (2λ_1_)^−1^, the extreme tail of the Satterthwaite approximation is exponential with rate (2*a*)^−1^. Since *a* < λ_1_ unless all the non-zero λ*_i_* are equal, the Satterthwaite approximation is increasingly anti-conservative in the extreme tail.

In our proposed approximation, increasing *k* by 1 takes the Satterthwaite remainder term 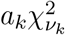, which is asymptotically lighter-tailed than the true distribution, and replaces it with a sum of two terms that has the correct asymptotic tail behavior. We can thus expect increasing *k* to improve the approximation for small *p*-values and large *m, n*, though not necessarily for large *p*-values or when *k* approaches min(*m*,*n*).

Supplementary Figure 11 shows how the accuracy of the Liu–Tang–Zhang, Satterthwaite, and leading-eigenvalue approximations compares across different *p*-values, using a single example with *N* = 5000. The advantage of the leading-eigenvalue approximation increases further out into the tail.

**Figure 11:**
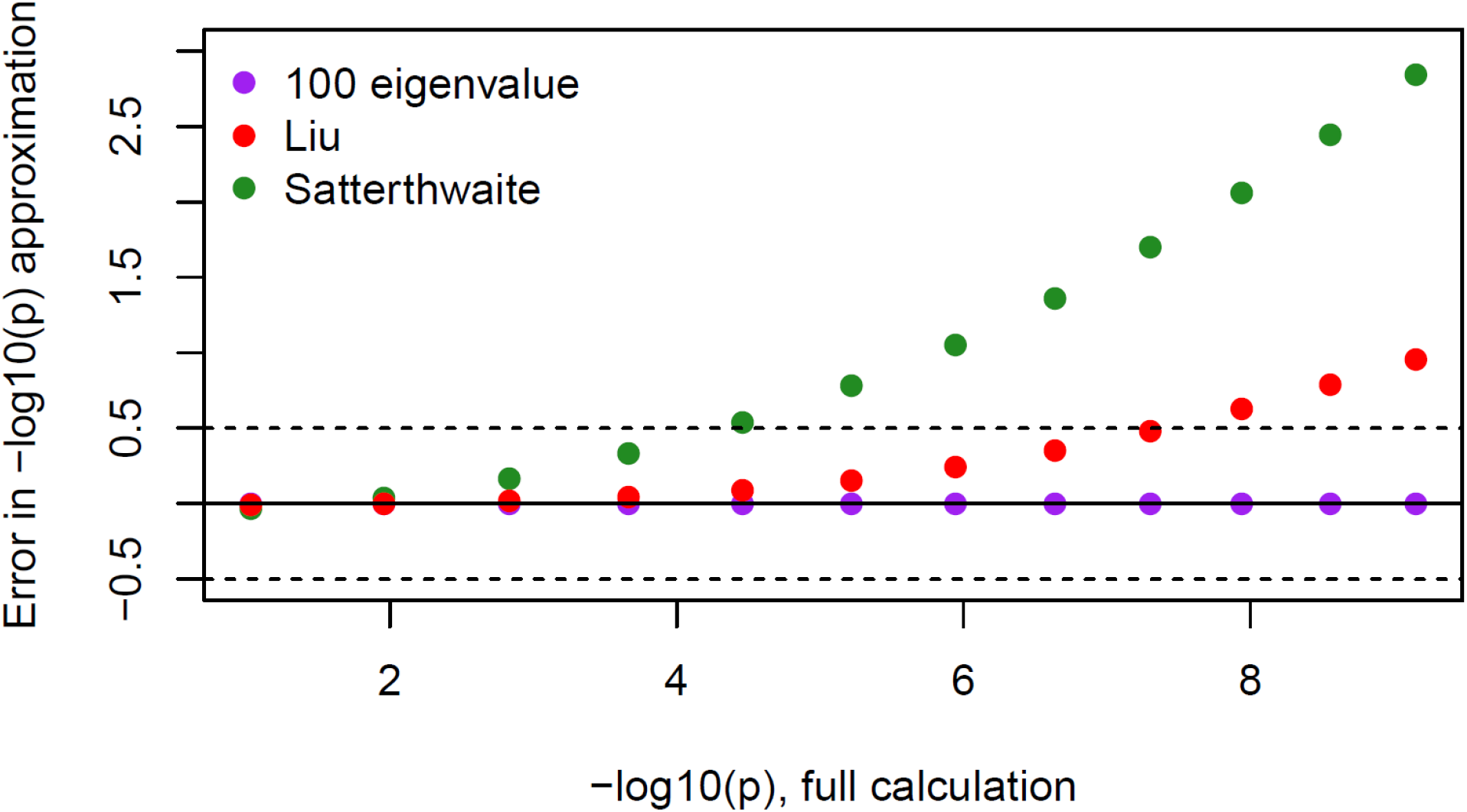
[Supplemental Figure] Dependence of accuracy on quantile, with *n* = 5000

## References

Anderson E., Bai Z., Bischof C., Blackford S., Dongarra J., Du Croz J., Greenbaum A., Hammarling S., McKenney A., and Sorensen D. (1999). LAPACK Users’ guide, volume 9. SIAM.

Berman S. M. et al. (1992). The tail of the convolution of densities and its application to a model of hiv-latency time. The Annals of Applied Probability, 2(2):481–502.

Blackford L. S., Petitet A., Pozo R., Remington K., Whaley R. C., Demmel J., Dongarra J., Duff I., Hammarling S., Henry G., et al. (2002). An updated set of basic linear algebra subprograms (blas). ACM Transactions on Mathematical Software, 28(2):135–151.

Chen G. K., Marjoram P., and Wall J. D. (2009). Fast and flexible simulation of dna sequence data. Genome research, 19(1):136–142.

Chen H., Lumley T., Brody J., Heard-Costa, N.L., Fox C. S., Cupples L. A., and Dupuis J. (2014). Sequence kernel association test for survival traits. Genetic epidemiology, 38(3):191–197.

Chen H., Meigs J. B., and Dupuis J. (2013a). Sequence kernel association test for quantitative traits in family samples. Genetic epidemiology, 37(2):196–204.

Chen H., Meigs J. B., and Dupuis J. (2013b). Sequence kernel association test for quantitative traits in family samples. Genetic epidemiology, 37(2):196–204.

Cirulli E. T. and Goldstein D. B. (2010). Uncovering the roles of rare variants in common disease through whole-genome sequencing. Nature Reviews Genetics, 11(6):415–425.

Davies R. B. (1980). Algorithm as 155: The distribution of a linear combination of χ 2 random variables. Journal of the Royal Statistical Society. Series C (Applied Statistics), 29(3):323–333.

Dixon J. R., Selvaraj S., Yue F., Kim A., Li Y., Shen Y., Hu M., Liu J. S., and Ren B. (2012). Topological domains in mammalian genomes identified by analysis of chromatin interactions. Nature, 485(7398):376–380.

Farebrother R. (1984). Algorithm as 204: the distribution of a positive linear combination of χ 2 random variables. Journal of the Royal Statistical Society. Series C (Applied Statistics), 33(3):332–339.

Fitzmaurice G. M., Laird N. M., and Ware J. H. (2012). Applied longitudinal analysis, volume 998. John Wiley & Sons.

Fuller W. (2011). Sampling Statistics. Wiley Series in Survey Methodology. John Wiley and Sons, Hoboken, NJ.

Galinsky K. J., Bhatia G., Loh P.-R., Georgiev S., Mukherjee S., Patterson N. J., and Price A. L. (2016). Fast principal-component analysis reveals convergent evolution of adh1b in europe and east asia. The American Journal of Human Genetics, 98(3):456–472.

Golub G. H. and Van Loan, C. F. (1996). Matrix computations, 3rd edition.

Goto K. and Van De Geijn, R. (2008). High-performance implementation of the level-3 blas. ACM Transactions on Mathematical Software (TOMS), 35(1):4.

Halko N., Martinsson P.-G., and Tropp J. A. (2011). Finding structure with randomness: Probabilistic algorithms for constructing approximate matrix decompositions. SIAM review, 53(2):217–288.

Hutchinson M. F. (1990). A stochastic estimator of the trace of the influence matrix for laplacian smoothing splines. Communications in Statistics-Simulation and Computation, 19(2):433–450.

Kang H. M., Sul J. H., Service S. K., Zaitlen N. A., Kong, S.-y., Freimer N. B., Sabatti C., Eskin E., et al. (2010). Variance component model to account for sample structure in genome-wide association studies. Nature genetics, 42(4):348–354.

Korobeynikov A., Larsen R. M., and Lawrence Berkeley National Laboratory (2016). svd: Interfaces to Various State-of-Art SVD and Eigensolvers. R package version 0.4.

Kuonen D. (1999). Saddlepoint approximations for distributions of quadratic forms in normal variables. Biometrika, 86(4):929–935.

Lee S., Abecasis G. R., Boehnke M., and Lin X. (2014). Rare-variant association analysis: study designs and statistical tests. The American Journal of Human Genetics, 95(1):5–23.

Lee S., Emond M. J., Bamshad M. J., Barnes K. C., Rieder M. J., Nickerson D. A., Team E. L. P., Christiani D. C., Wurfel M. M., Lin X., et al. (2012a). Optimal unified approach for rare-variant association testing with application to small-sample case-control whole-exome sequencing studies. The American Journal of Human Genetics, 91(2):224–237.

Lee S., with contributions from Larisa Miropolsky, and Wu, M. (2016). SKAT: SNP-Set (Sequence) Kernel Association Test. R package version 1.2.1.

Lee S., Wu M. C., and Lin X. (2012b). Optimal tests for rare variant effects in sequencing association studies. Biostatistics, 13(4):762–775.

Lin H., Wang M., Brody J. A., Bis J. C., Dupuis J., Lumley T., McKnight B., Rice K. M., Sitlani C. M., Reid J. G., et al. (2014). Strategies to design and analyze targeted sequencing data cohorts for heart and aging research in genomic epidemiology (charge) consortium targeted sequencing study. Circulation: Cardiovascular Genetics, 7(3):335–343.

Liu H., Tang Y., and Zhang H. H. (2009). A new chi-square approximation to the distribution of non-negative definite quadratic forms in non-central normal variables. Computational Statistics & Data Analysis, 53(4):853–856.

Loh P.-R., Tucker G., Bulik-Sullivan, B.K., Vilhjalmsson B. J., Finucane H. K., Salem R. M., Chasman D. I., Ridker P. M., Neale B. M., Berger B., et al. (2015). Efficient bayesian mixed-model analysis increases association power in large cohorts. Nature genetics, 47(3):284–290.

Lumley T. (2011). Complex surveys: a guide to analysis using R, volume 565. John Wiley & Sons.

Morrison A. C., Voorman A., Johnson A. D., Liu X., Yu J., Li A., Muzny D., Yu F., Rice K., Zhu C., et al. (2013). Whole genome sequence-based analysis of a model complex trait, high density lipoprotein cholesterol. Nature genetics, 45(8):899.

Price A. L., Patterson N. J., Plenge R. M., Weinblatt M. E., Shadick N. A., and Reich D. (2006). Principal components analysis corrects for stratification in genome-wide association studies. Nature genetics, 38(8):904–909.

Psaty B. M., O’Donnell C. J., Gudnason V., Lunetta K. L., Folsom A. R., Rotter J. I., Uitterlinden, A.G., Harris T. B., Witteman J. C., Boerwinkle E., et al. (2009). Cohorts for heart and aging research in genomic epidemiology (charge) consortium design of prospective meta-analyses of genome-wide association studies from 5 cohorts. Circulation: Cardiovascular Genetics, 2(1):73–80.

R Core Team (2016). R: A Language and Environment for Statistical Computing. R Foundation for Statistical Computing, Vienna, Austria.

Schifano E. D., Epstein M. P., Bielak L. F., Jhun M. A., Kardia S. L., Peyser P. A., and Lin X. (2012). Snp set association analysis for familial data. Genetic epidemiology, 36(8):797–810.

Sung Y. J., Korthauer K. D., Swartz M. D., and Engelman C. D. (2014). Methods for collapsing multiple rare variants in whole-genome sequence data. Genetic epidemiology, 38(S1):S13–S20.

Tropp J. A. (2011). Improved analysis of the subsampled randomized hadamard transform. Advances in Adaptive Data Analysis, 3(01n02):115–126.

Wu B., Pankow J. S., and Guan W. (2015). Sequence kernel association analysis of rare variant set based on the marginal regression model for binary traits. Genetic epidemiology, 39(6):399–405.

Wu M. C., Kraft P., Epstein M. P., Taylor D. M., Chanock S. J., Hunter D. J., and Lin X. (2010). Powerful snp-set analysis for case-control genome-wide association studies. The American Journal of Human Genetics, 86(6):929–942.

Wu M. C., Lee S., Cai T., Li Y., Boehnke M., and Lin X. (2011). Rare-variant association testing for sequencing data with the sequence kernel association test. The American Journal of Human Genetics, 89(1):82–93.

Yamazaki I., Bai Z., Simon H., Wang L.-W., and Wu K. (2010). Adaptive projection subspace dimension for the thick-restart lanczos method. ACM Transactions on Mathematical Software (TOMS), 37(3):27.

Yao L., Berman B. P., and Farnham P. J. (2015). Demystifying the secret mission of enhancers: linking distal regulatory elements to target genes. Critical reviews in biochemistry and molecular biology, 50(6):550–573.

